# Cerebral small vessel disease due to *Col4a1* mutations include cognitive impairment and white matter defects that can be modulated by targeting protein folding

**DOI:** 10.1101/2025.04.04.647207

**Authors:** Gaia Brezzo, Alexandra Pokhilko, Jon Moss, Juraj Koudelka, Maurits A. Jansen, Ross Lennen, Gerry Thompson, Nela Fialova, Sofia Marina Konstantinidou, Joyce Yau, Erin Boland, Anna Williams, Stuart M. Allan, Alessandra Granata, Sanjay Sinha, Tao Wang, Hugh S. Markus, Roman Fischer, M. Zameel Cader, Tom Van Agtmael, Karen Horsburgh

**Affiliations:** Institute for Neuroscience and Cardiovascular Research, University of Edinburgh, Edinburgh EH16 4SB, UK; School of Cardiovascular and Metabolic Health, University of Glasgow, University Avenue, Glasgow, G12 8QQ; UK Dementia Research Institute, University of Edinburgh, Edinburgh EH16 4SB, UK; Translational Molecular Neuroscience Group, Weatherall Institute of Molecular Medicine, Nuffield Department of Clinical Neurosciences, University of Oxford, Oxford, OX3 9DS, United Kingdom; Centre for Regenerative Medicine, Institute for Regeneration and Repair, The University of Edinburgh, Edinburgh EH16 4UU, UK; Division of Neuroscience, School of Biological Sciences, Faculty of Biology, Medicine and Health, The University of Manchester, Manchester, M13 9PT; Geoffrey Jefferson Brain Research Centre, Manchester Academic Health Science Centre, Northern Care Alliance NHS Foundation Trust, University of Manchester, Manchester, UK; Stroke Research Group, Department of Clinical Neurosciences Department, University of, Cambridge, Cambridge, UK; Cambridge Stem Cell Institute, Jeffrey Cheah Biomedical Centre, Puddicombe Way, Cambridge CB2 0AW; Division of Evolution, Infection and Genomics, School of Biological Sciences, Faculty of Biology, Medicine and Health, The University of Manchester, Manchester M13 9PT; Discovery Proteomics Facility, Target Discovery Institute, Nuffield Department of Medicine, University of Oxford, Oxford, OX3 7FZ, United Kingdom; Chinese Academy of Medical Sciences Oxford Institute, Nuffield Department of Medicine, University of Oxford, Oxford, UK

**Keywords:** Cerebral small vessel disease, Col4a1, white matter, proteomics, ER stress, basement membrane

## Abstract

White matter abnormalities are a hallmark of cerebral small vessel disease (cSVD) and are closely linked to cognitive decline and dementia. However, despite their clinical importance, underlying mechanisms remain poorly understood. Collagen IV, encoded by genes *COL4A1/COL4A2*, is a major component of the basement membrane, a specialised extracellular matrix (ECM) structure. Mutations in these genes cause a genetic form of cSVD. We tested the hypothesis that ECM defects caused by a *Col4a1* mutation lead to white matter pathology using an established mouse model of cSVD (*Col4a1^+/Svc^*). Behavioural testing with magnetic resonance diffusion tensor imaging, pathology and ultrastructural investigations of white matter were studied. The studies revealed that *Col4a1^+/Svc^* mice have cognitive impairments, reduced myelinating oligodendrocyte pools, axonal myelination defects, and altered white matter structural integrity. Proteomic analysis, of isolated white matter from *Col4a1^+/Svc^*mice, identified extensive changes to ECM composition and endoplasmic reticulum (ER) biology including ER stress induction. We also demonstrated that targeting protein folding to promote collagen secretion and reduce ER stress, increased myelinating oligodendrocytes and axon-glial integrity in *Col4a1^+/Svc^* mice. These data provide novel insight into the pathomolecular mechanisms of white matter abnormalities in cSVD and identify a modifiable pathway as a putative therapeutic target for cSVD.

## Introduction

Cerebral small vessel disease (cSVD), a disorder of the microvasculature, is a major cause of stroke, intracerebral haemorrhage (ICH), vascular cognitive impairment and dementia^1,2,3^. cSVD manifests on magnetic resonance imaging (MRI) as T2-white matter hyperintensities (WMH), lacunar infarcts, and cerebral microbleeds^1,4^. These features are shared in sporadic and familial (e.g. caused by *NOTCH3; HTRA1 and COL4A1/A2* mutations) forms of cSVD. WMH, which are a core feature of cSVD, become increasingly common in the general population with age and predict future stroke and dementia risk^5^. The pathogenic mechanisms resulting in white matter injury remain debated, but evidence supports endothelial cell dysfunction, compromised blood brain barrier integrity, perfusion alterations and microvascular inflammation as key instigators^6^. Most cases of cSVD are sporadic and associated with cardiovascular risk factors particularly hypertension, although genetic predisposition also plays a role^7^. Monogenic forms of cSVD, although rarer, are providing important insights into disease pathogenesis, with shared mechanisms between sporadic and monogenic disease being identified, particularly involving disruption of the neuro-glial-vascular unit and extracellular matrix (ECM)^8^.

One monogenic familial form of cSVD is caused by mutations in the genes *COL4A1 and COL4A2,* as part of COL4A1 syndrome (also referred to as Gould Syndrome)^9,10^. *COL4A1/A2* encode the protein collagen IV (COL4), an important component of basement membranes (BM), a specialised ECM structure^11^. cSVD due to *COL4A1/2* mutations is characterised by typical clinical and neuroimaging features of lacunar stroke and WMH, as well as intracerebral haemorrhage (ICH) and cerebral microbleeds on MRI^12^. However, the underlying pathology and mechanisms remain poorly defined. Although the familial syndrome is rare, its impact on cSVD in the population may be much greater. Typical familial mutations are much more common than expected in the general population, at about 1 per 1000, and are associated with stroke risk^2^. Moreover, rare coding variants that were predicted to be pathogenic have been detected in 2-3% of cSVD cases in population cohorts^13,14^, and common non-coding variants in *COL4A1/COL4A2* are risk factors for lacunar stroke^7^, ICH ^15^ and WMH ^16,17^. Thus, investigating COL4A1/2 syndrome can provide mechanistic insights into both sporadic and familial forms of cSVD.

To date research into cSVD due to *COL4A1/2* mutations has focused on ICH^11,12,13,18,19,20^, and no studies have investigated mechanistic links between COL4 and cerebral white matter abnormalities. Given that collagen IV is a key BM component we hypothesised that ECM defects culminate in white matter abnormalities. To address this we studied an established mouse model of cSVD in *Col4a1* mutant mice (*Col4a1^+/Svc^*) that harbours a glycine to aspartic acid mutation^21^, representative of ∼70% mutations found in COL4A1/2 Syndrome patients^22^. Using neuroimaging, pathology and ultrastructure investigations we establish that *Col4a1* mutations cause prominent white matter alterations including reduced myelination associated with loss of myelinating oligodendrocytes. Mechanistic proteomic investigations highlighted diverse protein alterations in the white matter with the endoplasmic reticulum (ER) being a key cellular compartment that is changed. Furthermore, targeting ER stress pharmacologically using the FDA-approved compound 4-phenylbutyric acid (PBA) conferred protection against white matter defects in COL4A1 disease, highlighting a putative therapeutic strategy.

## Methods

### Animals and PBA drug treatment

All animal procedures were performed in a UK Home Office–licensed facility in accordance with the Animal (Scientific Procedures) Act 1986 and the European Directive 2010/63/EU and approved by the Bioresearch and Veterinary Services Committee at University of Edinburgh and University of Glasgow. *Col4a1^+/Svc^* mice harbour a heterozygous glycine to aspartate point mutation (GD1064D) in exon 37 of *Col4a1* which affects collagen IV folding and secretion and recapitulates COL4A1/2 Syndrome^21,23,24^. *Col4a1^+/Svc^* mice are on a C57BL/6J background and wild-type (WT) littermates were used as controls. WT and *Col4a1^+/Svc^* mice were group housed on a 12-hour light/dark cycle with *ad libitum* access to food and water. Male and female mice were randomly allocated to experimental cohorts: longitudinal MRI at 3 and 8 months (*n*=8 WT, *n*=8 *Col4a1^+/Svc^*), pathology at 3 month (*n*=6 WT, *n*=6 *Col4a1^+/Svc^*); biochemical/proteomic analysis at 3-4m (*n*=7 WT, *n*=7 *Col4a1^+/Svc^*) and electron microscopy investigations at 3-4 months (*n*=3 WT, *n*=3 *Col4a1^+/Svc^*). Additional cohorts were used for behavioural analysis at 8 month old for nesting behaviour (*n*=6 WT, *n*=6 *Col4a1^+/Svc^*) and Y-maze tasks (*n*=5 WT, *n*=6 *Col4a1^+/Svc^*). *Col4a1^+/Svc^* mice were treated with 4-phenylbutyrate (PBA; 1g/kg/day, *n*=8) or vehicle (*n*=7)^24^ via drinking water from conception (by treating pregnant dams). Animals were randomly allocated to treatment/no-treatment group before genotyping and development of phenotypes. Experimenters were blind to genotype status and drug treatment.

### Brain imaging of white matter and blood flow

WT and *Col4a1^+/Svc^* mice underwent longitudinal imaging at 3 and 8 months of age to assess structural (T2-weighted), white matter microstructure (diffusion tensor imaging (DTI) and cerebral blood flow (arterial spin labelling, ASL) changes as described in the supplementary methods. Experimental animals were anesthetized under 5% isoflurane in oxygen for induction then placed in an MRI compatible holder (Rapid Biomedical, Wurzburg, Germany). Isoflurane was maintained at approximately 1.5% in oxygen during scanning. Rectal temperature and respiration were monitored and controlled throughout. Structural T2-weighted and DTI MRI data were collected using a 7T horizontal bore Biospec AVANCE preclinical imaging system with a 86umm quadrature volume coil and a two-channel phased-array mouse brain coil. ASL was performed using a Look-Locker FAIR single gradient echo (LLFAIRGE) sequence. DTI Fractional anisotropy (FA) maps were generated in Matlab. ASL maps were generated using the Bruker software (Paravision 360). For FA and ASL measures slices corresponding to 0.14mm – -0.66mm from Bregma and -1.82mm – -2.62mm from Bregma were processed in ImageJ for region of interest analysis. At the end of imaging brains were processed for pathological investigations.

### Immunohistochemical and ultrastructural investigation of white matter abnormalities

Mice were sacrificed under deep anaesthesia by transcardiac perfusion. PBA/vehicle treated mouse brains were collected without saline perfusion. Brains were post-fixed in 4% paraformaldehyde (PFA) for 24 h and embedded in paraffin wax. 6uµm thick coronal tissue sections were collected at corresponding MRI level coordinates (0.14mm and -1.82mm from Bregma). Tissue was sectioned using a rotary microtome (Leica Biosystems, Germany).

Sections were used for immunostaining using standard procedures. Briefly, sections were deparaffinised, blocked with 10% normal serum and 0.5% BSA for 1 h at room temperature before overnight primary antibody incubation at 4 °C. Sections were incubated with biotinylated secondary antibodies for 1 h at room temperature and either amplified with Vector ABC Elite Kit (Vector Labs, UK), before visualisation with 3,3′ diaminobenzidine tetrahydrochloride (DAB, Vector Labs, UK) or incubated with streptavidin (Invitrogen, UK). Myelin related abnormalities were assessed using SMI94 (BioLegend, 836504 1:15000) and MAG (Abcam, ab89780 1:10000) immunolabelling. CC1 (Merck, OP80 1:20) immunolabelling was used as a marker of mature oligodendrocytes, and CD31 (R&D, AF3628 1:100) used to assess endothelial cell density. Co-labelling of PDGRFα (R&D, AF1062 1:200) and Ki67 (Abcam, ab16667 1:100) was carried out to assess proliferation of oligodendrocyte precursor cells (OPC).

Transmission electron microscopy (EM) was performed to study ultrastructural white matter alterations. Mice were transcardially perfused with saline then fixative (4% paraformaldehyde, w/v, Sigma; 0.5% glutaraldehyde, v/v; Electron Microscopy Sciences, EMS; in 0.1 M phosphate buffer, PB, Sigma), and brains were post-fixed in the same fixative overnight. 50 µm coronal brain sections were collected using a vibratome, post-fixed with 1% osmium tetroxide (v/v in 0.1 M PB, EMS) and dehydrated in a series of ethanol dilutions (50%, 70%, 95% and 100%; Sigma) then acetone. Sections were then lifted into resin (Durcupan ACM, Fluka, Sigma), left overnight at room temperature, placed on greased microscope slides, coverslipped, and the resin cured at 65°C for 3 days. Regions of interest containing corpus callosum were cut from the slides, mounted on plastic blocks (Ted Pella) and ultrathin sections were taken (70 nm thick) for EM. Ultrathin sections were collected on to meshed copper grids, contrasted with uranyl acetate and lead citrate, and imaged with a JEOL transmission electron microscope (TEM-1400 plus). EM analysis was analysed as previously published^25^. Briefly, measures of axonal diameter, myelin thickness and inner tongue thickness were calculated from a measured area based on the assumption of circularity: diameter = 2 × √(area/π). 100 axons per animal were analysed using Fiji/Image J (Fiji.sc). Inner tongue thickness was calculated by subtracting the axonal diameter from the diameter of the innermost compact layer. G-ratios were calculated by dividing axonal diameter by axon, inner tongue and myelin area. Percentage of myelinated axons was calculated by dividing total number of axons by number of myelinated axons. Further qualitative analysis of abnormal phenotypes was carried out by counting axons with unfolding or unravelling of myelin and expressed as a percentage by dividing the total number of axons by total number of unfolding/unravelling axons.

### Behavioural assessments

#### Nesting behaviour

A nesting behaviour task was used to measure executive and cognitive function. One day prior to testing mice were singularly housed to assess individual nest building. Mice received one Nestlet (a pressed 2×2 inch square cotton material weighing 3g) one hour before the dark cycle. The following morning each individual nest was assessed on (1) appearance (Nestlet score), (2) height of the nest (in cm) and (3) percentage of shredded Nestlet. Scoring followed published criteria^26^.

#### Y-maze

Spatial working memory and spatial recognition memory were tested in a Y-maze with extra-maze visual cues to aid navigation^27^. The Y-maze apparatus consisted of three identical enclosed arms (50cm long x 11cm wide × 10cm high) made of black acrylic. The arms were mounted in a Y-shape, set at an angle of 120° from each other, with visual cues were placed at the end of each arm. For spatial working memory (spontaneous alternation) mice were placed in the centre of the Y-maze and allowed to explore all three arms for 5umin. The number of arm entries and the sequence of arms entered were recorded. The percentage alternation was calculated as number of alternations (entries into three different arms consecutively) divided by the total possible alternations (i.e. number of arms entered minus 2) and multiplied by 100. The number of errors the mice made were calculated as a re-entry to an arm just visited. The distance covered was also measured as an indirect measure of mobility. For spatial recognition memory, mice were placed into one of the arms of the maze (start arm) and allowed to explore the maze for 5umin with the entrance of one arm blocked off (training trial). After a 1-minute inter-trial interval (ITI), mice were returned to the start arm of the maze for the test phase with all three arms available for exploration (for a duration of 2 min) including the novel (previously blocked-off) arm. Spatial memory retention was measured as time spent in the novel arm calculated as a percentage of the total time spent in all three arms. Values were compared with the chance level for time exploring the three arms (33%).

### Biochemical and proteomic investigation of white matter enriched samples

#### Tissue processing

Mice were sacrificed under deep anaesthesia by transcardiac perfusion with PBS, whole brains were cut in the sagittal plane in three 2 cm slices and the corpus callosum was carefully dissected from adjacent grey matter. Following isolation of the corpus callosum, samples were homogenised in 1:10 (vol:vol) ice-cold RIPA buffer using a glass hand-held loose fit dounce homogeniser with 25 strokes. Samples were sonicated for 5 s and maintained in constant agitation for 2 hours at 4 °C. Finally, samples were centrifuged at 5000g for 2 min and the supernatant collected.

#### Proteomics

Up to 20 µg of white matter homogenate were prepared for LC-MS analysis using a 96-well S-Trap plate (Protifi LCC) following the manufacturer’s protocol. Samples were reduced with 5 µl 200 mM DTT and alkylated with 20 µl 200 mM IAM for 30 min each, acidified with 12% phosphoric acid 10:1 (v/v) sample to acid, transferred into the S-Trap well and precipitated with 7 parts 90% methanol in 100 mM TEAB to 1 part sample (v/v). Samples were washed 3x by spinning the S-trap plate at 1500xg for 30 s and the last step for 1 min, each time with fresh 90% methanol in 100 mM TEAB. The sample was resuspended in 50 µL 50 mM TEAB and digested with 0.8 µg trypsin (Promega) overnight at 37°C. Peptides were eluted from the S-Trap by spinning for 1 min at 1500xg with 80 µL 50 mM ammonium bicarbonate, 80 µL 0.1% FA and finally 80 µL 50% ACN, 0.1% FA. The eluates were dried down in a vacuum centrifuge and resuspended in loading buffer (2% ACN, 0.1% TFA) prior to high pH prefractionation and mass spectrometry acquisition.

#### High pH prefractionation on Agilent Bravo Assaymap

RP-S cartridges were primed with 100 µL ACN at 300 µL/min, equilibrated with 50 µL loading buffer (2% ACN, 0.1% TFA) at 10 µL/min followed by loading 120 µL sample at 5 µL/min. the cartridge cup was washed with 50 µL and an internal cartridge wash was performed with 25 µL loading buffer at 5 µL/min. Eight subsequent elution steps were done with buffer A (Water, pH 10) and buffer B (90% ACN, pH 10) at the following percentages of buffer B: 5%, 10%, 12.5%, 15%, 20%, 22.5%, 25% and 50%. The eluates of the following steps were directly concatenated 1+5, 2+6, 3+7 and 4+8. Fractions were dried down in a vacuum centrifuge and resuspended in loading buffer.

#### LC-MS/MS data acquisition

Peptides were analysed by LC-MS/MS as described^28^. Briefly, peptides were separated on a nEASY column (PepMAP C18, 75 µm x 500mm, 2 µm particle, Thermo) using a linear gradient of 2%-35% Acetonitrile in 5% DMSO/0.1% Formic acid at 250nl/min. MS data was acquired on a Q-Exactive mass spectrometer in DDA mode. MS1 resolution was set to 70,000 with an AGC target of 3E6. The top 15 most abundant precursors were selected for MS/MS with a resolution of 17,500, AGC target of 6E3 and an isolation window of 1.6 m/z. Fragments were generated with a normalized collision energy of 28% and selected precursors were excluded for 27 s.

#### Data analysis and protein identification and quantification

The proteomics analysis was based on protein identifications by MaxQuant. The raw data files were searched against either the reviewed Uniprot homo sapiens databased (retrieved 20180131) or mus musculus (retrieved 20190304) using MaxQuant^29^ version 1.6.10.43 and its built-in contaminant database using tryptic specificity and allowing two missed cleavages. The results were filtered to a 1% FDR.

More than 2000 proteins were identified in each mouse white matter sample with at least two unique peptides, except one outlier WT sample that only had 411 proteins. This sample was removed from downstream analysis. Differential protein analysis was done using 2595 proteins identified in both groups (WT and *Col4a1^+/Svc^*) in at least three biological replicates. Intensity values were normalized (divided) to median values in each sample to correct for between-sample differences, and missing values were replaced by median intensities for each protein in both groups^30,31^. Analysis of the differences in protein levels between groups was done with limma package in R using empirical Bayes method for two group comparison with the t-test of the eb.fit function^32^. The results were visualized on volcano plots with significance level *p* ≤ 0.05 and fold changes ≥ 1.5. All proteins from the differential analysis were combined in a single Excel sheet with all relevant information, including the annotation and location scores from String database^33^.

Pathway and protein-protein interactions (PPI) network analysis was done using all proteins identified during the differential analysis, including proteins exclusively present in only one group in ≥ three replicates, and present at zero levels in all samples of another group. Pathway network analysis was done using ClueGO plug-in^34^ in Cytoscape^35^ with KEGG and GO BP (Biological processes) databases. Only significantly enriched pathways were retained on the network (FDR<=0.05). The PPI network was constructed using

StringApp in Cytoscape, retaining only high confidence interactions, and proteins disconnected from the main network were omitted (except those present in the significantly enriched pathways from the ClueGO analysis). The location scores predicted by String database were used to colour the proteins according to their potential location based on the following prioritisation criteria: 1) proteins with high ER, endosome or golgi location scores (between 3 and 5=max) were assigned into group 1; 2) proteins with high mitochondria location score, but low score for the group 1 were assigned to group 2; 3) proteins with high cytosol or cytoskeleton location scores, which were not present in groups 1 and 2 were assigned to group 3; 4) proteins with high extracellular score, but low for the above groups were assigned to group 4. Groups 5 and 6 consisted of proteins with high plasma membrane or nuclear scores respectively, which were not present in the above groups.

#### Statistical analysis

MRI data (FA, ASL) were analysed by two-way mixed measures ANOVA with age and genotype as main factors with post-hoc Bonferroni. Immunohistochemistry, immunofluorescent and behavioural data were analysed by two-way ANOVA with post-hoc Tukey or Bonferroni. MAG grading was analysed using Kruskal-Wallis with post-hoc Dunn’s test or Mann Whitney-U for PBA-treatment. EM and PBA-treatment CC1 data was analysed with t-test. All statistical analysis was performed using Graphpad Prism (v10, Graphpad Software Inc.) or SPSS (v23, IBM Corporation). *p*<0.05 was considered to be statistically significant.

## Results

### Col4a1^+/Svc^ mice show widespread white matter alterations

To identify potential white matter defects, longitudinal fractional anisotropy (FA) measures from MR/DTI were taken from 3 and 8 months of old *Col4a1^+/Svc^* and WT mice (Figure 1A). White matter changes were assessed at two anatomical levels of the corpus callosum (rostral CC: Bregma 0.14 mm and caudal CC: Bregma -1.82 mm) and the internal capsule (IC, Bregma -1.82 mm) (Figure 1A). Two-way mixed measures ANOVA revealed significant effects of genotype (rostral CC: *F*(1,14)=8.14, *p*=.013; caudal CC: *F*(1,14)=13.3, *p*=.003; and IC: *F*(1,14)=20.85, *p*<.001) and age (rostral CC: *F*(1,14)=46,33, *p*<.001), as well as significant interactions (CC caudal: *F*(1,14)=5.18, *p*=.039; IC: *F*(1,14)=7.51, *p*=.016) (Figure 1B; Supplementary Table 1).

**Figure 1.**
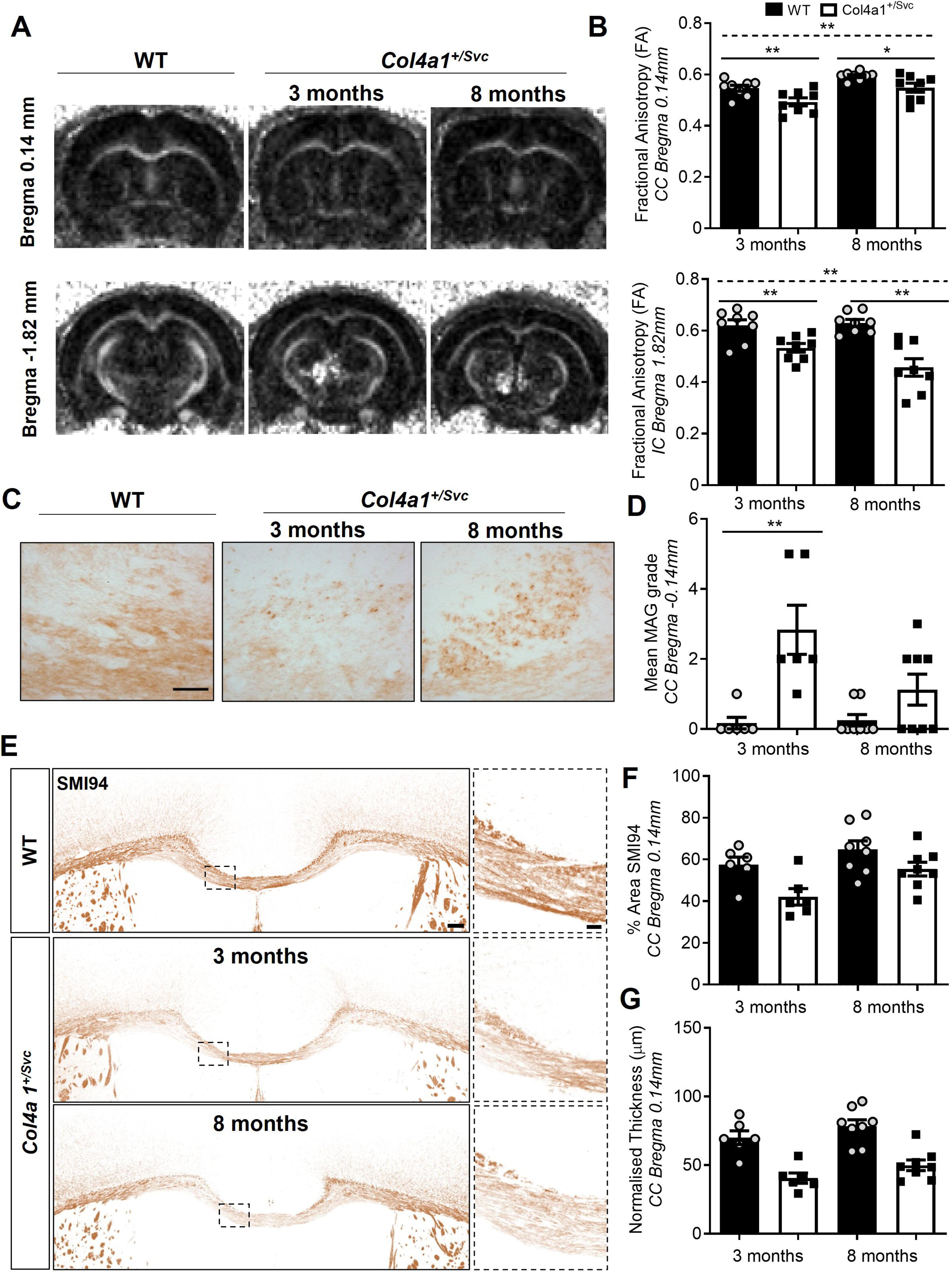
White matter abnormalities in Col4a1+/Svc mice. (A) Representative MR-DTI (FA) images from wild type (WT) and *Col4a1^+/Svc^* mice at 3 and 8 months of age. (B) Quantification of FA values in corpus callosum (CC) and internal capsule (IC) shows *Col4a1^+/Svc^* have significantly lower FA values compared to WT-littermates. (C) Representative images of MAG staining (indicative of axon-glial damage) in CC from WT and *Col4a1^+/Svc^* mice at 3 and 8 month of age. (D) Semi-quantitative grading in CC (Bregma 0.14mm) of MAG staining reveals accumulation of myelin debris and vacuolation, suggesting disruption of axon-glial integrity in *Col4a1^+/Svc^* mice. (E) Representative images of SMI94 staining in corpus callosum (Bregma level 0.14mm) from WT and *Col4a1^+/Svc^* animals at 3 and 8 month of age (F) Assessment of SMI94 % area coverage reveals decreased SMI94 density in *Col4a1^+/Svc^* compared to WT mice. (G) Normalised CC thickness by brain area. Quantification of CC thickness (Bregma level 0.14mm) indicates thinning of corpus callosum in *Col4a1^+/Svc^* mice at both ages. Scale bars (C) 50 μm (E) 100um; 20 μm (insert). In all panels, data are presented as mean ± SD and analysed by two-way mixed measures ANOVA with post hoc Bonferroni correction (B), by Mann Whitney U (D) or two-way ANOVA (F,G). *n*=6-8 \**p*<0.05, \*\**p*<0.01

To further define if these white matter defects were due to myelin and/or axonal pathology, we performed immunohistochemical investigations. Immunostaining for MAG, present at the axon-glia interface, revealed widespread accumulation of MAG^+^ myelin debris, across rostral (3mo: *p*=.0043) and caudal CC (3mo: *p*=.0206; 8mo: *p=*.0043) and IC (3mo: *p*=.0043; 8mo: *p=*.0115) in *Col4a1^+/Svc^* mice (Figure 1C-D; Supplementary Figure 1). Immunostaining for SMI94 (Figure 1E-F), a myelin basic protein marker, revealed myelin defects within the rostral CC in *Col4a1^+/Svc^* mice compared to WT. Two-way ANOVA analysis indicated significant effects of genotype (*F*(1,24) = 10.74, *p*=0.003) and age (*F*(1,24) = 7.318, *p*=0.012), but no interaction (*F*(1, 24) = 0.631, *p*=0.435). Assessment of callosal thickness across WT and *Col4a1^+/Svc^* mice showed white matter atrophy in *Col4a1^+/Svc^* mice (*F* (1,24) = 10.74, *p*<.001) but no significant effect of age (*F*(1,24) = 4.06, *p*=0.055), or interaction (*F*(1, 24) = 0.16, *p*=0.902) (Figure 1G). These data support that white matter defects due to a *Col4a1* mutation on MR-DTI can be explained at least partially by myelin loss and axon/glial damage.

### Early ultrastructural white matter abnormalities in Col4a1^+/Svc^ mice

To further define if white matter defects detected on MR-DTI were due to axonal/myelin pathology, we performed ultrastructural investigations of WT and *Col4a1^+/Svc^* mice (Figure 2A). Quantification of myelinated axons (Figure 2B) indicated a significant reduction in both axonal diameter (*p*=0.034) and myelin thickness (*p*=0.021) in *Col4a1^+/Svc^* mice (Figure 2B). However, there was no change in the inner tongue thickness (*p*=0.22) or G-ratio of myelinated axons (*p*=0.51). Overall, *Col4a1^+/Svc^*mice had 40% fewer myelinated axons (*p*=0.005) (Figure 2C), and showed more abnormal phenotypes (Figure 2D) including blebbing, outfoldings of myelin or unravelling of myelin sheaths compared to WT mice (*p*=0.002). These results support our MRI and histopathological observations of both myelin and axonal pathology in *Col4a1^+/Svc^*mice.

**Figure 2.**
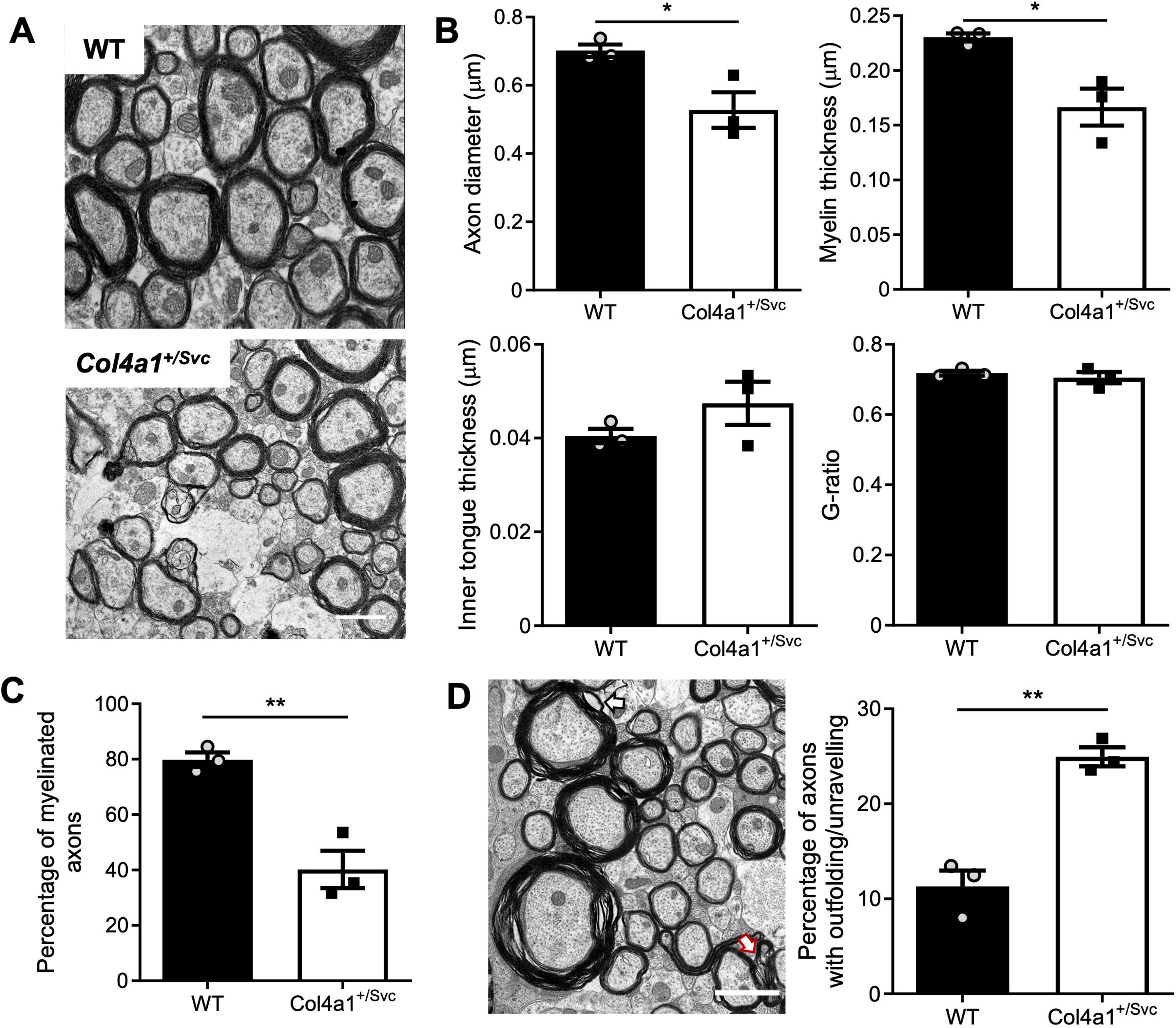
Ultrastructural evidence of axonal and myelin defects. (A) Representative images of myelinated and unmyelinated axons in CC (Bregma level 0.14mm) from WT and *Col4a1^+/Svc^* animals (n=3). (B) Quantification of axon diameter, myelin thickness, inner tongue thickness and G-ratio. *Col4a1^+/Svc^* animals have significantly smaller axons with decreased myelin thickness. (C) Quantification of myelinated vs unmyelinated axons in WT and *Col4a1^+/Svc^* mice. *Col4a1^+/Svc^* animals have significantly fewer myelinated axons compared to WT. (D) Abnormal phenotypes were identified as outfoldings (white arrow) or unravelling of myelin sheaths (red arrow). Quantification of these indicated that *Col4a1^+/Svc^* animals have a significantly higher percentage of outfoldings/unravelling compared to WT. In all panels, data are presented as mean ± SD and analysed by t-test. *n*=3, \**p*<0.05, \*\**p*<0.01. Scale bar 1 μm (A,D).

### Mature myelinating oligodendrocytes are reduced in Col4a1^+/Svc^ mice

Mature myelinating oligodendrocytes (OLs) are critical for the production, maintenance and regulation of axon myelination, enabling efficient neuronal communication maintaining cognitive function. Furthermore, as myelin is continuously remodelled in the central nervous system, turnover of mature OLs, from oligodendrocyte precursor cells (OPCs) is vital. Given the importance of both mature OLs and OPCs in maintaining efficient myelination, we investigated OLs and OPC integrity.

CC1 staining was used to determine the numbers of mature myelinating OLs in the CC of Col4a1^+/Svc^ mice and WT. We observed reduced numbers of mature OLs in Col4a1^+/Svc^ mice with significant effects of genotype (*F*(1,24)=15.05 *p*<0.001) and age (*F*(1,24)=4.896, *p*=0.037) but no interaction (*F*(1,24)=0.86, *p*=0.772) (Figure 3A-B). Reduced numbers of mature OLs could be a result of impaired proliferation, impaired differentiation and/or apoptosis. To unravel underlying mechanisms, we investigated the numbers and proliferation capacity of OPCs. We found a modest reduction in numbers of OPCs (stained by PDGFRα) in *Col4a1^+/Svc^* mice compared to WT. There was a significant effect of genotype (*F*(1,24)=5.205, *p*=0.032), but no significant effect of age (*F*(1,24)=2.847, *p*=0.104) or interaction (*F*(1,24)=1.986, *p*=0.172) (Figure 3C-D). Using Ki67 as a proliferative marker to quantify numbers of proliferating OPCs (defined as PDGFRa+Ki67+ cells), we observed increased proliferation in 3 month old *Col4a1^+/Svc^* mice (Figure 3C,E). Two-way ANOVA results indicate significant genotype (*F*(1,24)=5.505, *p*=0.028), age (*F*(1,24)=5.29, *p*=.030) and interaction effects (*F*(1,24)=8.018, *p*=0.009). This indicates proliferation capacity is not impaired in OPCs and does not account for the decreased numbers of mature OLs in *Col4a1^+/Svc^* mice. On the contrary, at least in young *Col4a1^+/Svc^* animals, OPC proliferation is increased, potentially to compensate for the loss of mature OLs caused by the *Col4a1* mutation.

**Figure 3.**
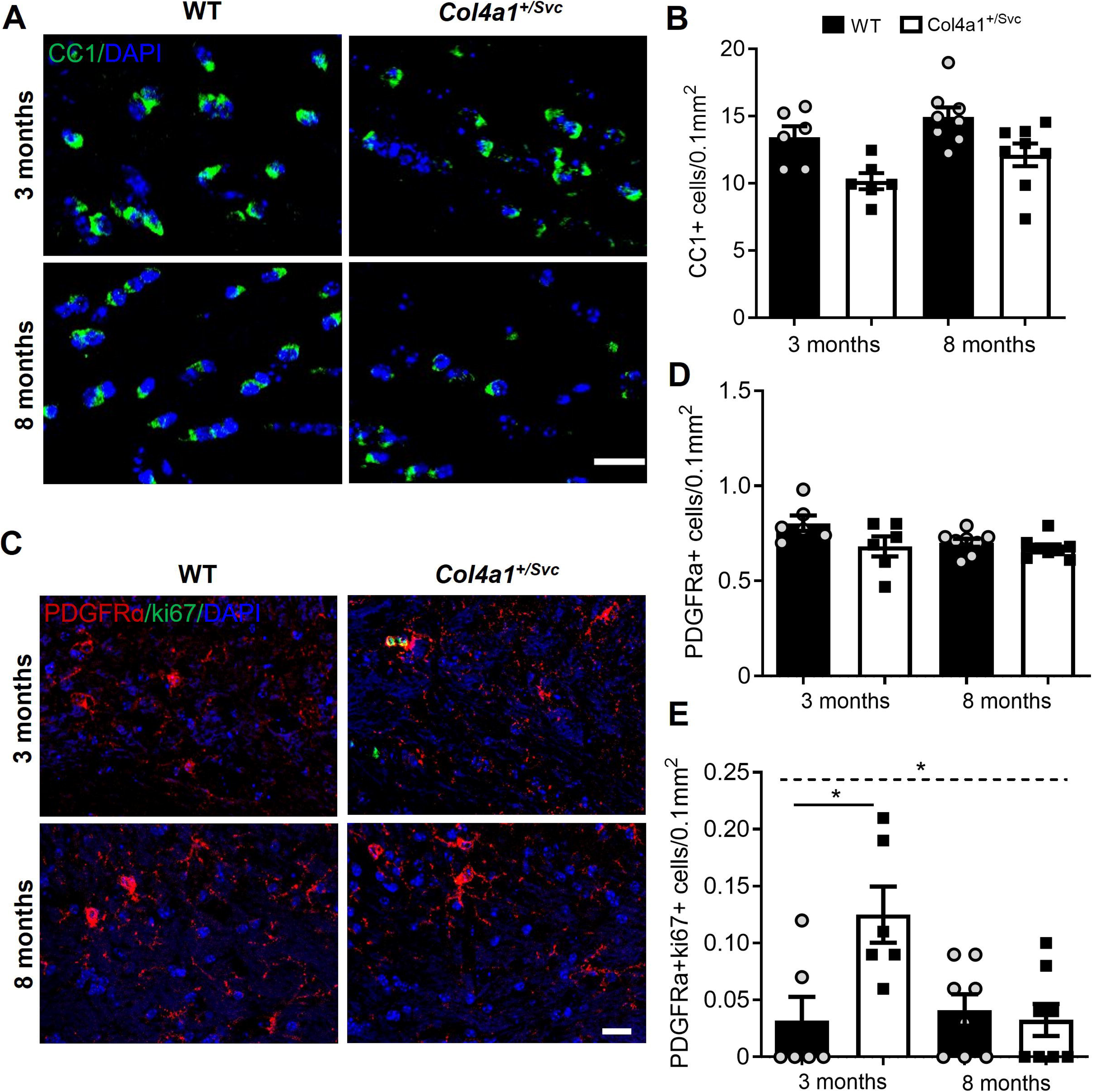
Mature myelinating oligodendrocytes are reduced in *Col4a1^+/Svc^* mice. (A) Representative images of CC1+ staining of mature oligodendrocytes in the corpus callosum (CC) (Bregma level 0.14mm) in *Col4a1^+/Svc^* and WT animals, at 3 and 8 month of age. (B) Quantification of number of mature oligodendrocytes (CC1+) reveals decreased oligodendrocyte numbers in *Col4a1^+/Svc^* compared to WT mice at 3 and 8 months of age. (C) Representative images of PDGFRu staining for oligodendrocyte precursor cells (OPC) and Ki67 staining for proliferating cells in CC (Bregma level 0.14mm) from WT and *Col4a1^+/S^*^vc^ animals at 3 and 8 months of age. (D) Quantification of the number of OPC (PDGFRu+) in 3 and 8 months animals indicates a modest decrease in OPCs at 3 months of age, but no differences across genotypes at 8 months of age. (E) Quantification of number of proliferating OPCs (PDGFRu+/Ki67+) in 3 and 8 month mice showed increased proliferation of OPCs in 3 month *Col4a1^+/Svc^* compared to WT mice. In all panels, data are presented as mean ± SD and analysed by two-way ANOVA \**p*<0.05. Scale bar 20 μm.

### Endothelial cell density and CBF alterations in white matter in Col4a1^+/Svc^ mice

Endothelial cell (EC) function and density can be reduced in cSVD and thus we wanted to determine if these are altered by the *Col4a1* mutation in white matter. EC density, identified as CD31+ immunostaining, was measured in rostral CC corresponding to white matter changes detected previously. We observed increased CD31 area coverage in *Col4^+/Svc^* animals compared to WT littermates. Two-way ANOVA revealed significant effects of genotype (*F(*1,24)=26.42, *p*<0.001) and age (*F*(1,24)=4.93 *p*=0.036), but no interaction (*F*(1,24)=0.786, *p*=0.384).

To determine if alterations in EC density in rostral CC may be linked to altered cerebral perfusion, we measured by ASL in *Col4a1^+/Svc^*compared to WT mice. Two-way mixed measures ANOVA results indicate significant genotype (*F*(1,14)=20.44, *p*<0.001), age (*F*(1,14)-7.46, *p*=0.016) and interaction effects (*F*(1,14)-10.82, *p*=0.005) (Figure 4D). We observed increases in CBF in *Col4a1^+/Svc^* animals compared to WT littermates at 3 months of age (*p*=0.001) but not 8 months. There is a significant decrease in CBF between *Col4a1^+/Svc^* at 3 and 8 months of age (*p*=0.003). Collectively these data, demonstrate age-dependent EC and CBF alterations in white matter of *Col4a1^+/Svc^* mice.

**Figure 4.**
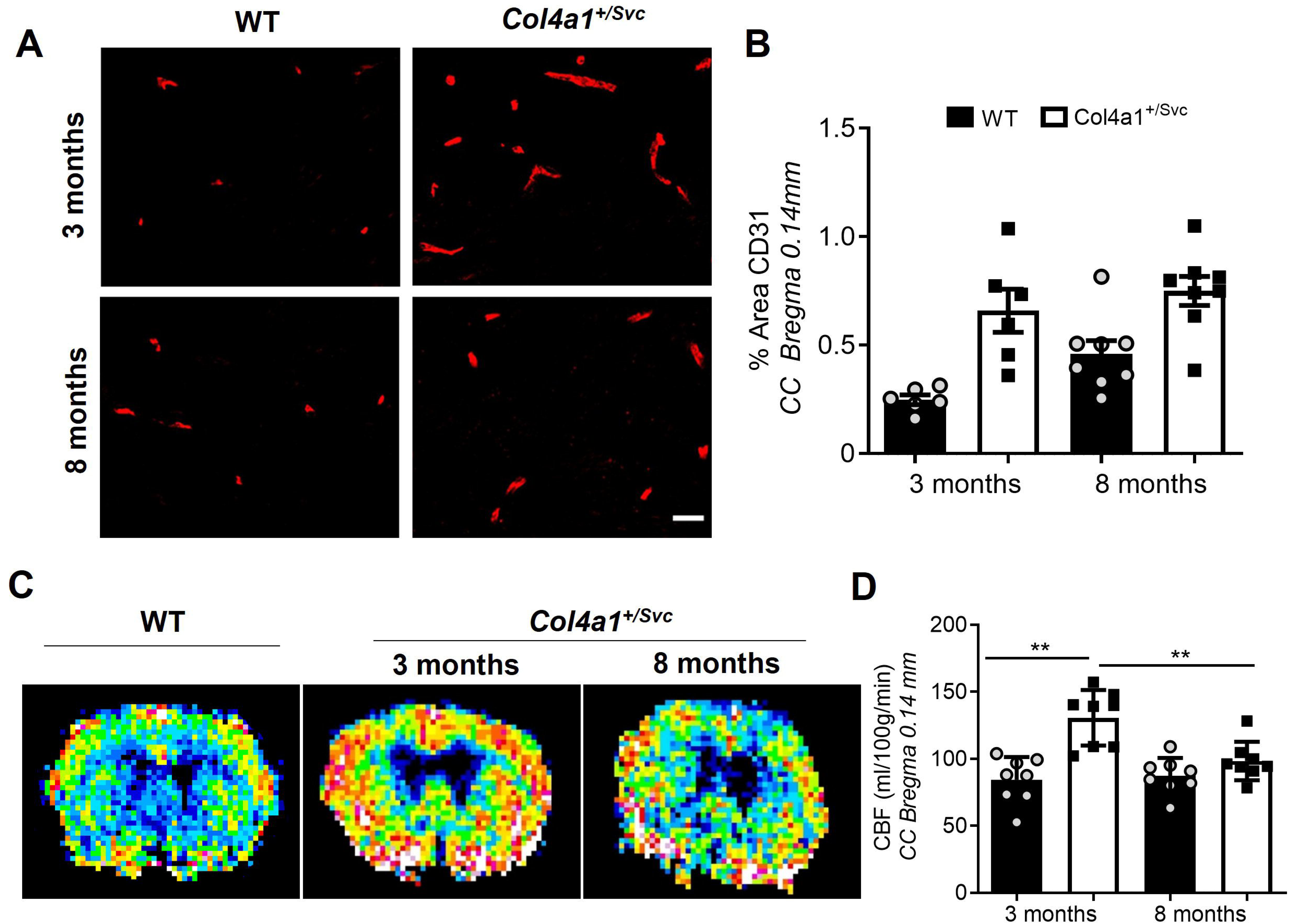
Increased endothelial cell density and perfusion in corpus callosum in *Col4a1^+/Svc^* mice. (A, B) Assessment of percentage area CD31 reveals increased CD31 density in *Col4a1^+/Svc^* mice compared to WT littermates (C) Representative MR Arterial Spin Labelling (ASL) images from WT and *Col4a1^+/Svc^* animals at Bregma level 0.14mm-0.66mm at 3 and 8 months of age. (D) Quantification of ASL values across genotype and age show that *Col4a1^+/Svc^* mice have regional differences in CBF. Notably in some regions like the CC, there is a significantly higher perfusion values compared to WT-littermates at 3 and 8 months of age. In all panels, data are presented as mean ± SD and analysed by two-way ANOVA (A, B) or two-way mixed measures ANOVA with post hoc Bonferroni correction \*\**p*<0.01. Scale bar 20 μm.

### Col4a1^+/Svc^ mice display impairment of cognition and executive function

Given the profound alterations in white matter integrity in *Col4a1^+/Svc^ mice*, we next investigated the impact of these changes on behavioural and cognitive functions at 8 months of age (Figure 5). Behaviour was firstly assessed with a nesting task (Figure 5A) which we have previously showed to be a sensitive marker for brain executive function^37^. Nest appearance was significantly lower for *Col4a1^+/Svc^* mice (*p*=0.012, Figure 5B). We next assessed spatial working and spatial recognition memory with the Y-maze (Figure 5C). We found no significant difference in the percentage of spontaneous alternation of arm entries between WT and *Col4a1^+/Svc^* mice (*p*=0.36) (Figure 5D). Further analysis of the percentage of errors (as an index of short-term working memory) revealed a non-significant (*p*=0.061) trend towards more errors in the *Col4a1^+/Svc^* mice (Figure 5E). To check for differences in locomotor activity, we measured distance travelled between *Col4a1^+/Svc^* mice and WT littermates and found no differences, (*p*=0.126) indicating intact locomotor activity (Figure 5F).

**Figure 5.**
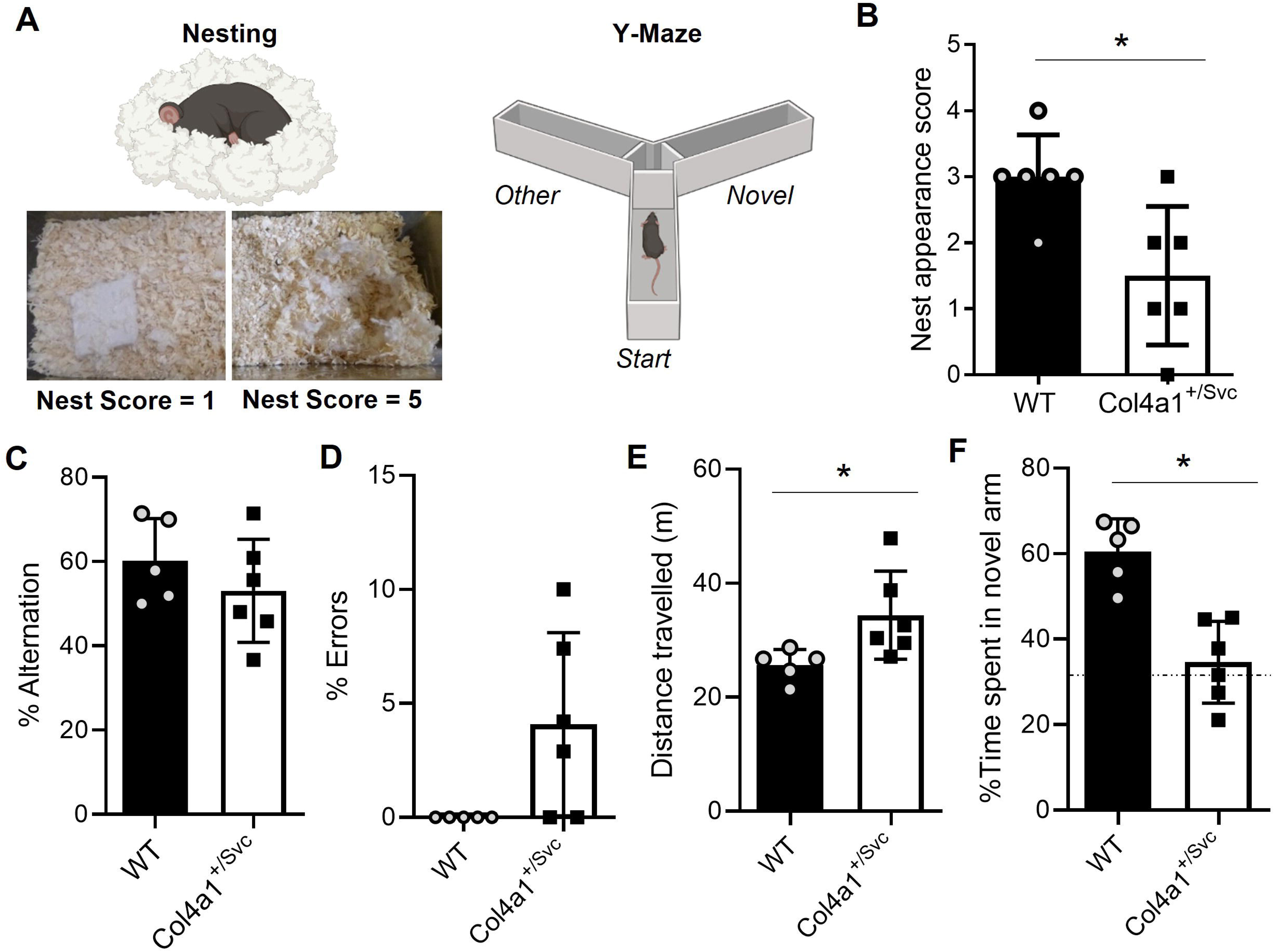
Cognitive and executive brain function is impaired in *Col4a1+/Svc* mice. (A) Schematics of Nesting and Y-Maze apparatuses. Nesting ability was scored on appearance of nests and representative images of nests with score of 1 and 5 respectively are shown. (B) Quantification of nesting ability (in *Col4a1+/Svc* animals and WT littermates at 8 months of age. Col4a1+/Svc mutants show decreased nesting ability, indicative of impaired executive function. (C) Percent spontaneous alternation between 3 arms of Y-maze in WT and *Col4a1+/Svc* mice were not significantly different. (D) Percent errors during spontaneous alternation in WT and *Col4a1+/Svc* mice indicates *Col4a1+/Svc* tend to make more errors than WT mice. (E) Total distance travelled during the spontaneous alternation task by WT and *Col4a1+/ Svc* mice. *Col4a1+/Svc* mice travelled significantly greater distance overall compared to the WT. (F) Percent time WT and *Col4a1+/Svc* mice spent exploring the novel arm during retention phase were significantly different with *Col4a1+/Svc* mice spending less time exploring the novel arm compared to the WT. WT/*Col4a1+/Svc*; n=5/6, Mann-Whitney U tests were used for statistical analyses. Measures are presented as a mean ±SD. * *p*<0.05). Dotted line represents chance level.

To assess short-term spatial recognition memory, a two-trial (acquisition and retention) Y-maze test with a 1 min inter trial interval (ITI) was used (Figure 5C). In the acquisition phase, one of the three arms is blocked off, allowing mice to explore two arms only (e.g. ‘start’ and ‘other’ arm), We found no significant difference in the time spent in the ‘start’ (*p*=0.942) or ‘other’ (*p*=0.991) arm between WT and *Col4a1^+/Svc^* mice. However, following a 1 min ITI were all three arms of the Y-maze are available for exploration, *Col4a1^+/Svc^*mice spent a significantly lower percentage of time exploring the novel arm compared to WT mice (*p*=0.004) (Figure 5G). Collectively, these studies indicate that *Col4a1* mutation affects behavioural measures including cognitive and executive function.

### Proteomic investigation highlights ER stress and ECM defects

To gain insight into the underlying molecular changes of the white matter abnormalities in *Col4a1^+/Svc^* mice, we combined mass spectrometry with off-line high-pH reversed-phase fractionation to maximise proteome coverage. Proteomics analyses revealed 142 differentially expressed proteins between WT and *Col4a1^+/Svc^*animals (*p*≤ 0.05, fold change ≥ 1.5; Figure 6A), in addition, 34 proteins were identified exclusively in *Col4a1^+/Svc^* mice, resulting in a total of 176 differentially expressed proteins (Supplementary Table 3).

**Figure 6.**
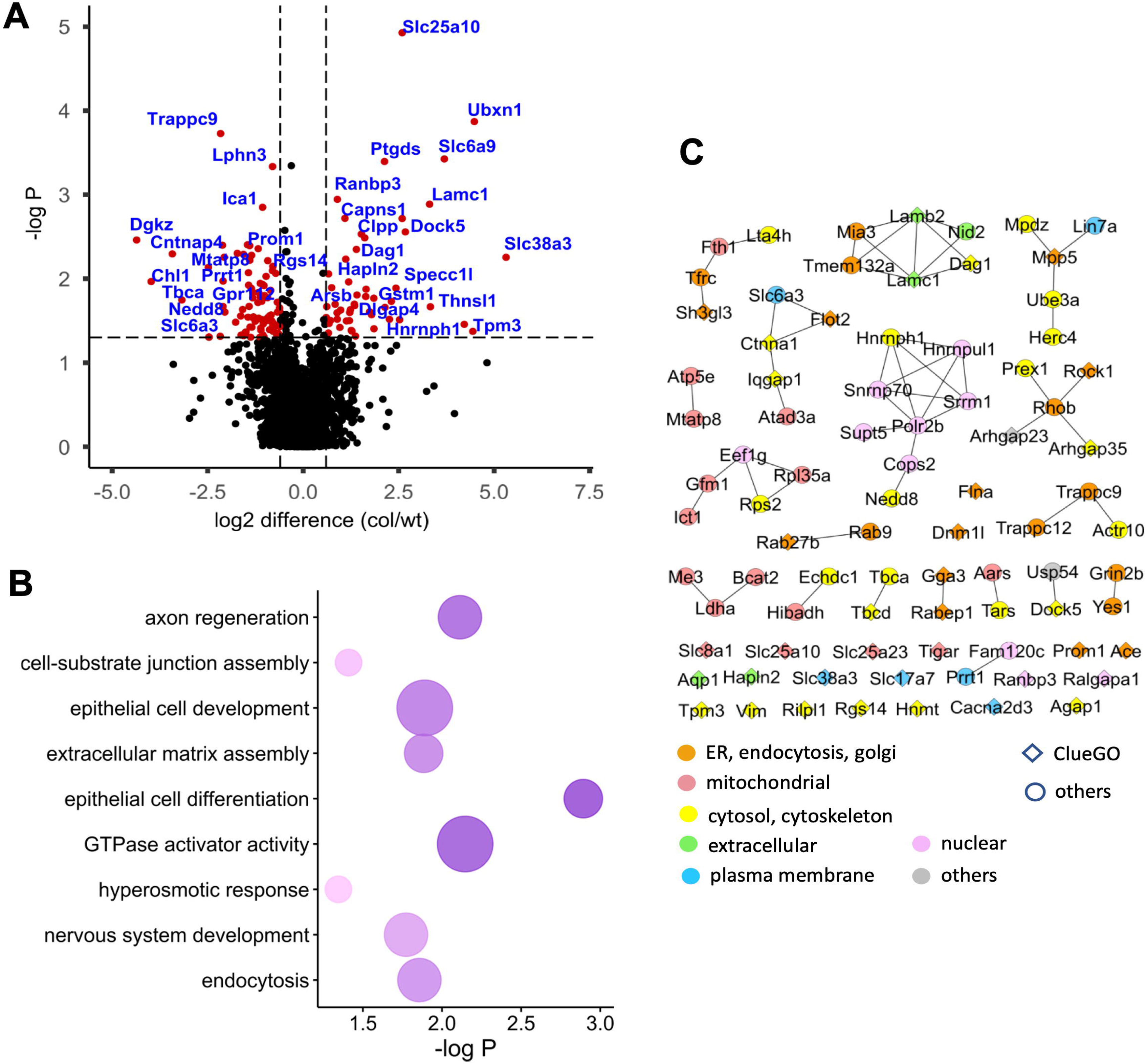
Proteomic investigations highlight white matter proteins and pathways/networks altered in *Col4a1^+/Svc^* mice. (A) Volcano plot of differentially regulated proteins between WT and *Col4a1^+/Svc^* mice, based on all proteins identified in at least 3 biological replicates in each group. Proteins that are present at significantly different levels (*p* ≤ 0.05) are marked red. (B) Biological pathways, significantly associated with *Col4a1* mutation (FDR ≤ 0.05), using KEGG and GO BP databases. The analysis was done on differentially regulated proteins from A, using ClueGO plug-in in Cytoscape. Darker colours of the pathway terms correspond to higher significance. (C) Network of interactions of differentially regulated proteins between WT and *Col4a1^+/Svc^* mice. The networks were built using the STRING database with a high confidence of interactions. The symbol colours indicate potential location of the protein, predicted based on String location score as described in Methods. The shape of the symbols shows whether the protein is also present in the pathway terms from ClueGO analysis.

To explore the biological function of these differentially expressed proteins, we performed pathway analysis with ClueGO plug-in in Cytoscape, using KEGG and GO BP databases. ECM and axon-related terms were significantly represented amongst the differentially regulated pathways (Figure 6B, Supplementary Table 3). In addition, epithelial cell differentiation, endocytosis, GTPases activity and ionic/osmotic changes were significantly represented. To further explore the potential location and inter-relationships between the differentially expressed proteins, we constructed a String PPI network. This analysis showed localisation of these proteins to the ER and mitochondria, as well as to the ECM and plasma membrane (Figure 6C). We observed altered levels of laminin and nidogen, two key BM components indicating compositional white matter BM defects. Furthermore, many of the differentially expressed proteins were related to the ER/mitochondrial stress responses (e.g., Ubxn1, Tigar, Dnm1l, Slc6a9, Supplementary Table 3), which we previously showed can occur in other tissues in *Col4a1^+/Svc^* mice^23, 24^.

### Targeting protein folding partially ameliorates white matter abnormalities in Col4a1^+/Svc^ mice

As the ER was a key cellular compartment altered with the *Col4a1* mutation and many differentially expressed proteins were related to ER stress, we wanted to determine the extent to which targeting ER stress and promoting collagen IV secretion could exert white matter protection. *Col4a1^+/Svc^* mice were treated orally (from conception) with the FDA-approved compound, PBA which promotes protein trafficking and reduces ER stress^19, 24^, and compared to untreated *Col4a1^+/Svc^* mice.

We firstly undertook MAG immunostaining to assess axon-glial integrity (Figure 7A), which we previously report to be affected in *Col4a1^+/Svc^* mice. Notably, PBA treatment reduced the extent of axon-glial breakdown compared to untreated *Col4a1^+/Svc^* mice (*p*=0.034) providing evidence that improving protein folding is beneficial in this disease context. We next set out to ascertain if PBA treatment also ameliorated the deficiency in myelinating OLs (CC1+ cells) numbers (Figure 7B). We indeed show that PBA treatment significantly increased the number of OLs in *Col4a1^+/Svc^* mice (*p*=0.028). Thus, the improved axon-glial integrity in PBA treated *Col4a1^+/Svc^*mice may be attributed to increased numbers of myelinating OLs. These data support a beneficial effect of PBA treatment and targeting of ER stress on white matter defects in cSVD due to a *Col4a1* mutation.

**Figure 7.**
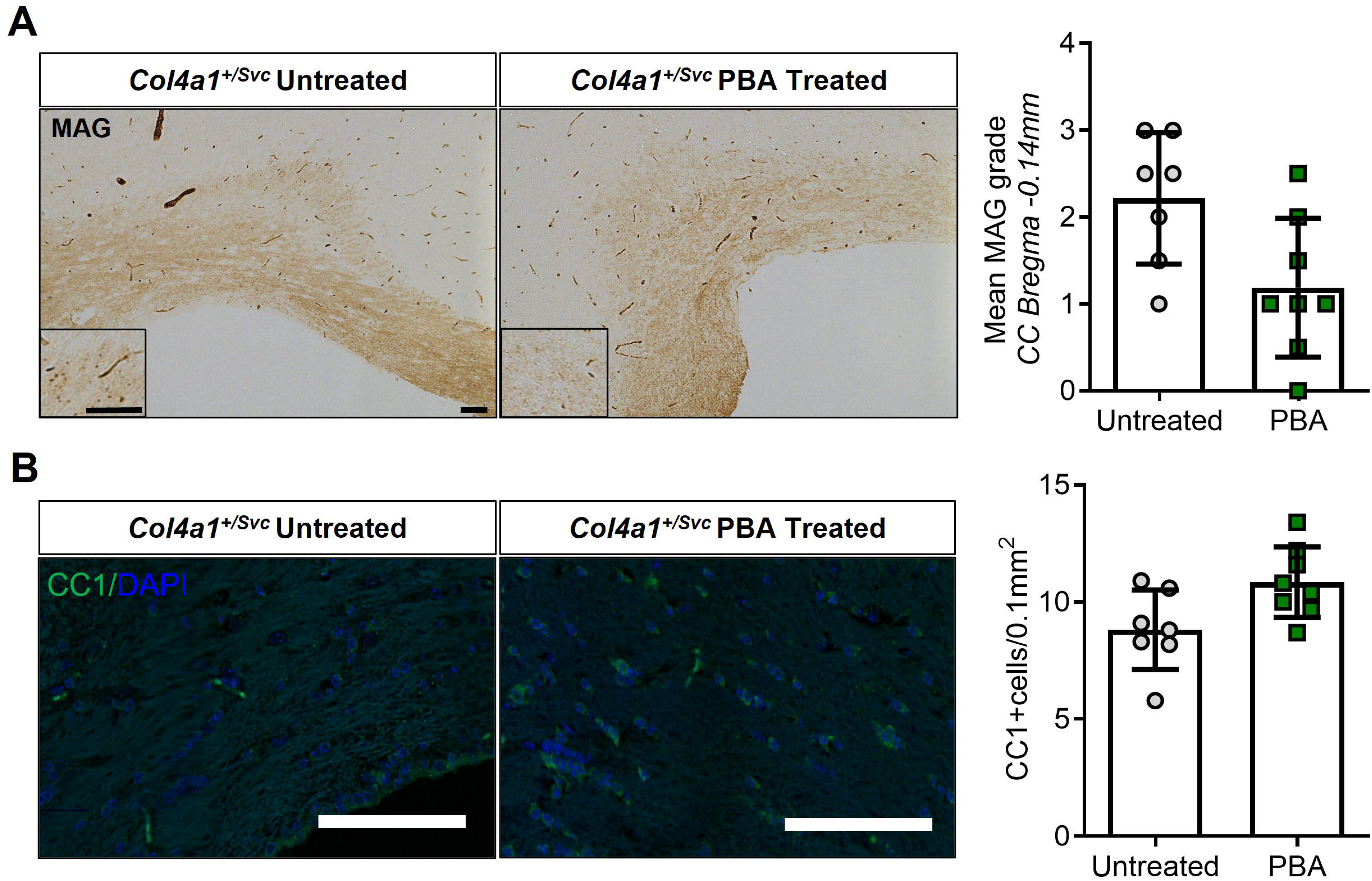
PBA treatment promotes white matter integrity and myelinating oligodendrocytes in *Col4a1^+/Svc^* mice. (A) Representative images of MAG immunostaining in corpus callosum in PBA-treated and untreated *Col4a1^+/Svc^* mice. (Scale bars= 50μm). Grading of myelin debris accumulation highlights a significant accumulation of myelin debris in untreated *Col4a1^+/Svc^* mice compared to PBA-treated *Col4a1^+/Svc^* mice. Data analysed by Mann Whitney U. (B) Representative images of CC1 staining of mature oligodendrocytes in corpus callosum in *Col4a1^+/Svc^* animals. (Scale bar= 100 μm). Quantification of number of mature oligodendrocytes (CC1+ve) reveals increased oligodendrocyte numbers in PBA treated *Col4a1^+/Svc^* compared to untreated mice (untreated *Col4a1^+/Svc^* n=7; PBA treated *Col4a1^+/Svc^* n=8). Data analysed by t-test. All data are represented as mean ± SD \**p*<0.05.

## Discussion

White matter abnormalities are a prominent feature of familial and sporadic cSVD and are associated with increased risk of cognitive impairment and dementia. Here, we demonstrate the presence of white matter defects and impaired cognitive function in a well-established mouse model of genetic cSVD. We established that these defects are associated with extensive changes to ER and ECM proteins, suggesting that ECM defects lead to white matter pathology. Importantly, we show that targeting ER protein folding using the FDA-approved chemical chaperone PBA rescued white matter pathology, informing on a future putative treatment strategy for cSVD associated with collagen IV variants.

We firstly utilised neuroimaging approaches (MR-DTI), which we previously determined can detect microstructural alterations in damaged white matter in the rodent brain ^42, 43^, and showed reduced FA in subcortical white matter tracts (corpus callosum, internal capsule) of *Col4a1^+/Svc^* mice. With immunohistochemical (SMI94, MAG) and ultrastructural EM investigations, we furthermore established that decreases in FA are a result of both myelin and axonal pathology and may be mediated by loss of mature OLs.

Cerebral hypoperfusion is proposed as a central common mechanism underlying white matter abnormalities in cSVD. However, studies are often cross sectional in nature and CBF related mechanisms are not always investigated in relation to white matter integrity^44^. Using ASL to quantify regional alterations in CBF in white matter regions, we showed increased CBF at 3 months (and to a lesser extent at 8 months) of age in *Col4a1^+/Svc^* mice , which was associated with increased vascular density and remodelling. Similarly, small peripheral vessels in these mice show enhanced endothelial dependent vasodilation^18^. We hypothesise that this may reflect early disease mechanisms that precede later suppression/reduction of CBF. Although clinical studies that have investigated regional CBF alterations early in cSVD are lacking, cerebral hyperperfusion has been reported in asymptomatic APOE4 individuals with white matter disruption before the onset of dementia^46^. Indeed, we show impaired short-term spatial recognition memory, a key memory domain affected in cSVD patients^47^, in *Col4a1^+/Svc^* mice by 8 months of age. Similarly, in a milder *Col4a1* model (*Col4a1^+/G^*^394^*^V^*) impaired working memory was observed at 12 months of age and was attributed to impaired blood flow responses to neuronal stimulation, supporting a loss of vascular function in COL4A1 disease^45^.

Our proteomic approach, which significantly increases ECM protein coverage^48^, provided insight into underlying molecular changes within the white matter of *Col4a1^+/Svc^* mice. Our data revealed extensive changes in BM and ECM composition, with upregulation of laminin (Lamb2, Lamc1), Dag1, a dystroglycan protein involved in laminin assembly and cell survival, Nid2, a BM protein crucial for BM stabilisation, cell adhesion and implicated in smooth muscle cell contractability. We previously showed that endothelial cell defects drive peripheral vascular dysfunction in COL4A1 disease^18,56^ and again find significant changes in proteins involved in vascular integrity, BBB function, angiogenesis and EC migration (Ptgds, Flna, Dock5, Rock1)^57,58^.Taken together these data, demonstrate the impact of a Col4a1 mutation on wider BM composition with implications for vascular function.

*Col4a1* mutations affect the folding and secretion of the collagen IV protein, inducing ER stress,^11,23,24^ and proteostasis, mechanisms which have been implicated in neurodegeneration^55^. We show that proteins critical for collagen secretion and folding (Tango1, Hsp47)^10^, and proteins involved in stress responses are altered in *Col4a1^+Svc^* mice, including those linked to ER stress (e.g. Ubxn1, Slc6a9 and Capns1) or linked to hypoxia/oxidative stress (e.g. Tagln2, Clpp, Aqp1, Slc25a10). We also observed, altered levels of stress-related proteins that are involved in axonal growth, (Usp47, Stmn2, Arhgap23) furthermore linking ECM dysfunction to axonal changes.

Finally, to determine if promoting collagen folding and secretion can rescue white matter defects in COL4A1 disease we treated *Col4a1^+/Svc^* mice with PBA. We and others have previously showed, that PBA treatment can lower ER stress and increase extracellular collagen IV levels^10,19,20,24^ and therefore hypothesised that PBA treatment would protect against BM and vascular remodelling ameliorating white matter pathology. Indeed, we observed increased OLs numbers and improved white matter integrity in *Col4a1^+/Svc^* mice (reduction of MAG debris) but treatment was not fully restorative. This could point to pathogenicity of mutant protein secretion as seen in other mouse tissues and/or models ^24, 59–61^ and the need to further modulate ECM-derived defects.

In conclusion, we established that *Col4a1* mutations cause prominent white matter alterations and cognitive changes, mediated by dysregulated ER biology and ECM composition, which can be targeted pharmacologically to ameliorate white matter pathology. These data provide novel insight into the diversity of pathomolecular mechanisms of white matter abnormalities in cSVD and identify a modifiable pathway as a putative therapeutic target for the treatment of cSVD-related white matter pathology.

## Acknowledgements

We would like to acknowledge the contributions of students Dan Sandler, Sara Castro Devesa, Siham Nimale, Oonagh Thin, Cameron Thomson (University of Edinburgh) who helped provide preliminary analysis of MRI, immunohistochemical and behavioural studies. The technical support provided by Steve Mitchell at the TEM facility, University of Edinburgh and Raphael Heilig at the Discovery Proteomics Facility, University of Oxford are gratefully appreciated.

## Authors Contribution Statement

K.H. directed and designed the research study. G.B. performed the neuroimaging studies with support from M.J, R.L. and G.T. G.B. J.M., A.W. were responsible for the ultrastructural studies and analysis. G.B. was responsible for the immunohistochemical studies and analysis. R.F. was responsible for LC-MS analysis. A.P. and L.H. undertook the downstream proteomic analysis. E.B performed intervention study under supervision of T.V.A.. G.B., A.P., L.H., K.H. and Z.C. analysed and interpreted the data with critical input from T.V.A, A.N., A.G., S.S., T.W., H.S.M. and S.A.N. provided critical reading of the manuscript. K.H. wrote the manuscript with support from G.B., A.P. and T.V.A with comments from all authors. All authors edited the manuscript and approved the final submission.

## Statements and Declarations

### Ethical Considerations

All animal protocols were performed in conformity with the Animal Research: Reporting of In Vivo Experiments (ARRIVE) guidelines. All animal experiments were conducted in accordance with the Animal (Scientific Procedures) Act 1986 and local ethical approval at the University of Edinburgh and University of Glasgow and were performed under personal and project licences granted by the Home Office.

### Consent for Publication

Not applicable.

### Data availability

Mass spectrometry proteomics data have been deposited to the ProteomeXchange Consortium via the PRIDE partner repository with the dataset identifier PXD057204. The processed data are supplied as Supplementary Tables.

Supplementary material for this paper can be found at http://jcbfm.sagepub.com/content/by/supplemental-data

