## Supplementary figures and images for "Cerebral small vessel disease due to *Col4a1* mutations include cognitive impairment and white matter defects that can be modulated by targeting protein folding"

### Supplemental Figure 1

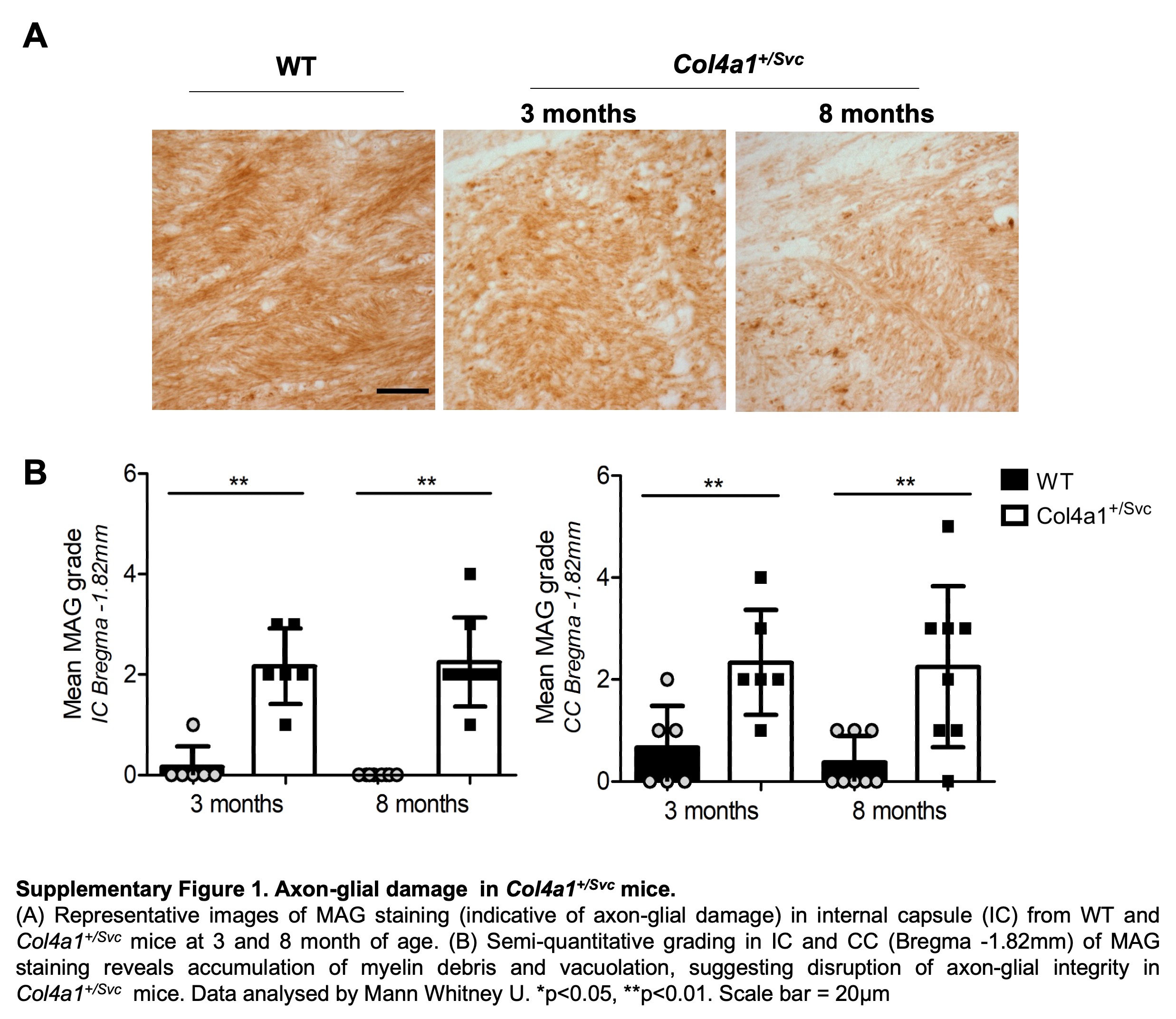
