## Supplemental Methods for "Cerebral small vessel disease due to *Col4a1* mutations include cognitive impairment and white matter defects that can be modulated by targeting protein folding"

**Supplementary Methods**

*Magnetic Resonance Imaging Parameters*

MRI was performed on a 7 T horizontal bore Biospec AVANCE neo preclinical imaging system equipped with a 116 mm bore gradient insert (Bruker BioSpin GmbH, Germany, maximum gradient strength 660 mT/m). Mice were anaesthetised with 1.5–2% isoflurane (Zoetis Ltd., London UK) in oxygen/air (50/50, 1 L/min) and secured in a cradle (Rapid Biomedical GmbH, Rimpar, Germany). The respiration rate and rectal temperature were monitored (Model 1030 monitoring and gating system, Small Animal Instruments Inc. Stony Brook, NY, USA), with body temperature maintained at 37°C by a heat fan. A 86 mm quadrature volume coil (Bruker BioSpin GmbH, Germany) was used for transmission with signal reception by a two-channel phased-array mouse brain coil (Rapid Biomedical GmbH, Rimpar, Germany).

Scout images were taken to confirm correct positioning and the magnetic field was optimised using automated 3D field mapping routine. For all subsequent sequences, the field of view was 19.2 × 19.2 mm and the slice thickness 0.8 mm. For anatomical imaging, 17 coronal slices covering the entire brain were acquired using a T2-weighted Rapid Acquisition with Relaxation Enhancement (RARE) sequence with the following parameters: matrix size 192 × 192, TR 2300 ms, effective TE 36 ms, RARE factor 4, number of signal averages 4. The scan time was 7 min 21 s.

The diffusion tensor (DT-)MRI protocol consisted of 4 T2-weighted and sets of diffusion-weighted (b=1000s/mm2) Echo Planar Imaging (EPI) volumes acquired with diffusion gradients applied in 30 non-collinear directions, producing a total of 34 volumes. The acquisition matrix was 128 × 128 and the number of shots was 4. The TR and TE times for each EPI volume were 2000 and 20 ms respectively and the scan time 4 min 32 s. DTI parameter maps were generated in Matlab.

Arterial Spin Labelling (ASL) was performed using a 2-shot flow alternating inversion recovery (FAIR) spin-echo EPI sequence. Thirteen images were acquired after a slice-selective or global adiabatic inversion pulse (TI: 15 – 7500 ms, interleaved), inversion slab thickness = 3.8 mm, allowing determination of T1. Total scan time was 9.27 min for a 96×96 imaging matrix. The flip angle was 90°. Two slices were obtained for each animal at each time point. From these images, T1sel and T1nonsel were calculated using a non-linear least square fit. CBF maps were generated using the Bruker software (Paravision 360) using equation [1] as discussed by Bell et al. (JMRI 1998; 8:2140-1245).

CBF=λ×60000 ×1/(T1,blood)×((T1,global)/(T1,sel) -1)	[1]

With the blood-tissue partition coefficient for water, λ = 90 mg/L, T1,blood = 2300ms and a unit conversion factor of 60000 ms/min.

Lastly, a susceptibility-weighted gradient echo sequence was acquired with TR 306 ms, TE 7 ms, flip angle 30°, a 192×192 matrix and 4 signal averages in 4 minutes.
