## Supplemental Table 1 for "Cerebral small vessel disease due to *Col4a1* mutations include cognitive impairment and white matter defects that can be modulated by targeting protein folding"

| **Region of Interest** | **WT 3mo** | **Col4^+/Svc^ 3mo** | **WT 8mo** | **Col4^+/Svc^ 8mo** | **Statistics** | **% Change** |
| --- | --- | --- | --- | --- | --- | --- |
| **Corpus Callosum**  Bregma 0.14 to  -0.66mm | 0.55 ± 0.032 | 0.49 ± 0.044 | 0.59 ± 0.016 | 0.55 ± 0.050 | **Genotype; *F*(1,14) = 8.14, *p*=0.013***  **Age; *F*(1,14) = 46.33, *p*<0.001***  Age*Genotype; *F*(1,14)=0.271 *p*=0.610 | 3mo:  -10.9%  8mo:  -6.80% |
| **Corpus Callosum**  Bregma  -1.82 to  -2.62mm | 0.45 ± 0.015 | 0.42 ± 0.04 | 0.48 ± 0.04 | 0.40 ± 0.041 | **Genotype; *F*(1,14) = 13.30, *p*=0.003***  Age; *F*(1,14) = 0.07, *p*=0.792  **Age*Genotype; *F*(1,14)=5.18 *p*=0.039*** | 3mo:  -6.70%  8mo:  -16.7% |
| **Internal Capsule**  Bregma  -1.82 to  -2.62mm | 0.62 ± 0.062 | 0.53 ± 0.048 | 0.63 ± 0.039 | 0.46 ± 0.096 | **Genotype; *F*(1,14) = 20.85, *p*<0.001***  Age; *F*(1,14) = 4.53, *p*=0.052  **Age*Genotype; *F*(1,14)=7.51 *p*=0.016*** | 3mo:  -14.5%  8mo:  -27.0% |

**Supplementary Table 1. Reduced FA in *Col4a1^+/Svc^* mice.**

Quantification of FA in white matter regions across genotype and age. *Col4a1^+/Svc^* have significantly lower FA values compared to WT-littermates at 3 months and 8 months of age. Data analysed by two-way mixed measures ANOVA with post-hoc Bonferroni correction. Bold font indicated significant main effect and/or significant interaction.
