## Supplemental Table 2 for "Cerebral small vessel disease due to *Col4a1* mutations include cognitive impairment and white matter defects that can be modulated by targeting protein folding"

| **Region of Interest** | **WT 3mo** | **Col4^+/Svc^ 3mo** | **WT 8mo** | **Col4^+/Svc^ 8mo** | **Statistics** | **% Change** |
| --- | --- | --- | --- | --- | --- | --- |
| **Corpus Callosum**  Bregma 0.14 to  -0.66mm | 84.36 ± 17.10 | 130.66 ± 20.69 | 87.36 ± 13.33 | 98.39 ± 14.31 | **Genotype; *F*(1,14) = 20.44, *p*<0.001***  **Age; *F*(1,14) = 7.46, *p=0*.016***  **Age*Genotype; *F*(1,14)=10.82 *p*=0.005*** | 3mo:  +54.9%  8mo:  +12.6% |
| **Corpus Callosum**  Bregma  -1.82 to  -2.62mm | 115.70 ± 15.34 | 156.17 ± 29.68 | 120.62 ± 28.14 | 114.58 ± 44.15 | Genotype; *F*(1,14) = 1.87, *p*=0.194  Age; *F*(1,14) = 4.09, *p*=0.063  **Age*Genotype; *F*(1,14)=6.57 *p*=0.023*** | 3mo:  +35%  8mo:  +5% |
| **Internal Capsule**  Bregma  -1.82 to  -2.62mm | 169.04 ± 23.50 | 241.98 ± 30.93 | 164.28 ± 20.52 | 244.03 ± 46.36 | **Genotype; *F*(1,14) = 33.83, *p*<0.001***  Age; *F*(1,14) = 0.22, *p*=0.883  Age*Genotype; *F*(1,14)=0.141 *p*=0.713 | 3mo:  +43.2%  8mo:  +48.6% |

**Supplementary Table 2. Increased cerebral perfusion in *Col4a1^+/Svc^* mice.**

Quantification of ASL values in white matter regions across genotype and age. *Col4a1^+/Svc^* have significantly higher perfusion values compared to WT-littermates at 3 months of age. Perfusion levels return to WT-baseline levels in corpus callosum, but remain elevated in the internal capsule at 8 months of age. Data analysed by two-way mixed measures ANOVA with post hoc Bonferroni correction. Bold font indicated significant main effect and/or significant interaction.
